# The nuclear sulfenome of *Arabidopsis*: spotlight on histone acetyltransferase GCN5 regulation through functional thiols

**DOI:** 10.1101/2024.04.24.590918

**Authors:** Barbara De Smet, Xi Yang, Zuzana Plskova, Carmen Castell, Alvaro Fernandez-Fernandez, Avilien Dard, Amna Mhamdi, Didier Vertommen, Kai Xun Chan, Sébastien Pyr dit Ruys, Joris Messens, Pavel I. Kerchev, Frank Van Breusegem

**Affiliations:** Department of Plant Biotechnology and Bioinformatics, Ghent University, 9052 Ghent, Belgium; Center for Plant Systems Biology, VIB, 9052 Ghent, Belgium; Structural Biology Brussels Laboratory, Vrije Universiteit Brussel, 1050 Brussels, Belgium; Structural Biology Research Center, VIB, 1050 Brussels, Belgium; Brussels Center for Redox Biology, 1050 Brussels, Belgium; de Duve Institute and MASSPROT platform, Université Catholique de Louvain, 1200 Brussels, Belgium; Research School of Biology, The Australian National University, Acton ACT 2601, Australia; Mendel University in Brno, Zemedelska, 613 00 Brno, Czech Republic

**Keywords:** *Arabidopsis*, GCN5, nucleus, oxidation, post-translational modification, sulfenome

## Abstract

Partial reduction of oxygen during energy generating metabolic processes in aerobic life forms results in the production of reactive oxygen species (ROS). In plants, ROS production is heightened during periods of both abiotic and biotic stress, which imposes a significant overload on the antioxidant systems. Hydrogen peroxide (H_2_O_2_) holds a central position in cellular redox homeostasis and signalling, playing an important role by oxidising crucial cysteines to sulfenic acid (-SOH), considered as a biologically relevant post-translational modification (PTM). Until now, the role of the nucleus in the cellular redox homeostasis has been relatively underexplored. The regulation of histone-modifying enzymes by oxidative PTMs on redox-active cysteines or tyrosine residues is particularly intriguing as it allows the integration of redox signalling mechanisms with chromatin control of transcriptional activity. One of the most extensively studied histone acetyltransferases is the conserved GENERAL CONTROL NONDEPRESSIBLE 5 (GCN5) complex. This study investigated the nuclear sulfenome in *Arabidopsis thaliana* by expressing a nuclear variant of the Yeast Activation Protein-1 (YAP) probe, identifying 225 potential redox-active nuclear proteins subject to sulfenylation. Mass spectrometry analysis further confirmed the sulfenylation of GCN5 at specific cysteine residues, with their functional significance and impact on the protein-protein interaction network assessed through cysteine-to-serine mutagenesis.

**Highlight:** Protein cysteine thiols are post-translationally modified under oxidative stress. Through the *in vivo* capturing of nuclear proteins undergoing sulfenylation in *Arabidopsis*, we highlight the functionality of particular cysteines in the histone acetyltransferase GCN5.

## Introduction

Partial reduction of oxygen during energy generating metabolic processes in aerobic life forms leads to production of reactive oxygen species (ROS) such as superoxide, hydrogen peroxide (H_2_O_2_), and hydroxyl radical. In plants, ROS production is particularly exacerbated during periods of abiotic and biotic stress which imposes a significant overload on the omnipresent antioxidant systems (Mittler *et al*., 2022; Mata-Pérez *et al*., 2023). Extended or severe acute stress episodes ultimately overcome the enzymatic and non-enzymatic antioxidant defences and culminate in oxidative damage to proteins, nucleotides, and lipids, leading to impaired plant growth and development. However, fine-tuned doses of ROS are also required for normal cellular functioning that relies on redox signalling mechanisms (Petrov and Breusegem, 2012; Noctor and Foyer, 2016). Unlike other ROS, H_2_O_2_ has a relatively long half-life (milliseconds to seconds) and high activation energy which makes it stable enough to diffuse and generate gradients from its source and react selectively with protein metal centres and specialised thiols (Waszczak *et al*., 2015; Willems *et al*., 2023). Its unique physico-chemical properties and highly regulated production and detoxification position H_2_O_2_ in the centre of the cellular redox homeostasis and signalling (Huang *et al*., 2024). The reaction of H_2_O_2_ with cysteine thiols first results in the formation of highly an unstable sulfenic acid (-SOH) which can be further oxidized to sulfinic (-SO_2_H) or sulfonic acid (-SO_3_H), or can react with other cysteine residues, forming disulfide bonds (Akter *et al*., 2018; Huang *et al*., 2021). Proteins that undergo sulfenylation are an integral part of redox networks, and they are linked by kinetically controlled redox switches within the proteome, where protein conformation, macromolecular interactions, trafficking through thiol disulfide protein structures, activity and function play an essential role. To comprehensively identify sulfenylated proteins, various redox proteomics workflows are utilised. These workflows employ chemical or proteinaceous probes to target sulfenylated cysteine residues and enrich them for identification using mass spectrometry. This approach allows for the proteome-wide analysis of cysteine S-sulfenylation, providing insights into the redox regulation of proteins. Recent advances in chemoproteomics and redox proteomic strategies have significantly improved the *in vivo* applicability of these techniques, enabling a better understanding of protein redox signalling (Willems *et al*., 2016; Huang *et al*., 2019). One of these approaches turned the Yeast Activation Protein-1 (YAP1) into an affinity reagent towards sulfenylated cysteines *in cellulo* (Takanishi and Wood, 2011). Expression of such a YAP1 probe in *Arabidopsis thaliana* plant cells, allowed the identification of nucleo-cytosolic and plastid proteins that underwent sulfenylation by H_2_O_2_ (Waszczak *et al*., 2014; De Smet *et al*., 2019; Wei *et al*., 2020). One of the advantages of the YAP1 probe approach over chemical probes is the ability to cargo it to subcellular locations and to trap sulfenylated proteins within a spatio-temporal context of interest.

The compartmentalised production and scavenging of ROS are instrumental for the successful integration of redox signalling in plant growth and developmental programs (Foreman *et al*., 2003; Manzano *et al*., 2014; Considine and Foyer, 2021). This results in fine-tuned doses of ROS that affect only the spatially and temporally accessible part of the proteome. In the plant cell, chloroplasts, mitochondria, and peroxisomes are the main centers of ROS production, but simultaneously are also equipped with a robust repertoire of enzymatic and non-enzymatic antioxidants. Other cellular compartments, such as the apoplast and cytosol, are also important players in the overall cellular redox homeostasis. They either primarily generate ROS or detoxify ROS released from other subcellular organelles, respectively. Intriguingly, the role of the nucleus in the cellular redox homeostasis has remained largely underexplored until now. Even though anecdotic evidence points towards certain nuclear localised ROS production mechanisms, the predominant ROS detected in the nucleus is likely entering via nuclear pores from the cytosol and chloroplasts (Foyer and Hanke, 2022). Moreover, physical association between chloroplasts and nuclei and formation of stromules (tubular protrusions stretched from the plastid body filled with stroma) have been observed under high light which likely facilitates the transfer of chloroplast-generated H_2_O_2_ into the nucleus (Exposito-Rodriguez *et al*. 2017). The major small-molecule antioxidants, ascorbate, glutathione, along with various antioxidant enzymes, are also present in the nucleus (Vivancos *et al*., 2010; Mittler, 2017). Interestingly, relocation of antioxidant enzymes from the cytosol to the nucleus can be observed under stress conditions, supporting the notion of active regulation of nuclear redox homeostasis (Tsang *et al*., 2014; Baker *et al*., 2023).

The epigenetic landscape consisting of histone marks, DNA methylation, and histone variants is dynamically rearranged under abiotic and biotic stress conditions (Ueda and Seki, 2020; Lloyd and Lister, 2022; Hannan Parker *et al*., 2022). This is achieved through epigenetic regulators that either deposit or remove histone marks and (de)methylate DNA (Candela-Ferre, 2024). The activity of histone modifying enzymes can be transcriptionally regulated but is also affected by post-translational modifications (Yang and Li, 2020; Zheng *et al*., 2023). Regulation of histone modifying enzymes by oxidative post-translational modifications on redox-active cysteine or tyrosine residues is particularly intriguing since it allows integration of redox signalling mechanisms with chromatin control of transcriptional activity (Plskova *et al*., 2024).

Histone acetylation facilitates the access of the transcriptional machinery to actively transcribed genome regions and is an important mechanism that accompanies numerous developmental and defence processes (Rymen *et al*., 2019; Barrero-Gil *et al*., 2022; Chen *et al*., 2024). Among the most widely studied histone acetyltransferases that deposit histone acetylation marks is the evolutionary conserved GENERAL CONTROL NONDEREPRESSIBLE 5 (GCN5), which is part of the transcriptional co-activator Spt–Ada– Gcn5 Acetyltransferase (SAGA) complex. The protein subunits of SAGA are organised in four distinct modules: histone acetylation (HAT), deubiquitination (DUB), SPT, and TAF module (Wu *et al*., 2021). The *Arabidopsis* GCN5 (*At3g54610*) preferentially acetylates lysine residues on Histone H3 (H3K9, H3K14, H3K27) but also several non-histone targets such as ADA2 are acetylated by GCN5 (Benhamed *et al*. 2006; Mao *et al*. 2006; Earley *et al*. 2007). Apart from the histone acetyltransferase (HAT) domain, GCN5 contains a bromodomain that is suggested to act as a reader of epigenetic information which recognizes acetylated lysine residues (Zeng and Zhou 2002; Li and Shogren-Knaak, 2009). Intriguingly, GCN5 is also part of the plant-specific PAGA complex which apart from GCN5 and ADA2A contains four plant-specific subunits (ING1, SPC, SDRL, and EAF6). PAGA and SAGA display an antagonistic effect that can lead to repression of gene expression while at the same time both complexes independently deposit moderate and high levels of histone acetylation marks, respectively, facilitating gene expression. In contrast to SAGA, which is implicated in both developmental and stress programs, PAGA targets specifically genes controlling plant morphology (Wu *et al*., 2023).

Plants lacking GCN5 are dwarfed and display various developmental defects, including decreased fertility, impaired floral development, and root meristem differentiation (Vlachonasios *et al*., 2003; Kim *et al*., 2009; Kornet and Scheres, 2009). Among the gene targets of GCN5 are important developmental players involved in root stem cell maintenance such as *WUSCHEL-RELATED HOMEOBOX5* (*WOX5*), S*HORT-ROOT* (*SHR*), and *SCARECROW* (*SCR*) and the floral regulators *AGAMOUS*, *WUSCHEL,* and *LEAFY* (Kim *et al*., 2018; Bertrand *et al*., 2003; Cohen *et al*., 2009). Many stress-regulated genes are also targeted by GCN5 positioning this histone modifier as a master regulator at the interplay between developmental and stress programs (Kim *et al*., 2020). Upon heat stress, GCN5 is actively recruited to the chromatin and deposits H3K9Ac and H3K14Ac marks to the promoter regions of the crucial heat stress regulators *HEAT SHOCK TRANSCRIPTION FACTOR A3* (*HSFA3*) and *UV-HYPERSENSITIVE 6* (*UVH6*). In the absence of GCN5, the heat induction of *HSFA3* and *UVH6* is impaired which correlates with increased heat sensitivity (Hu *et al*. 2015). Wheat mutants lacking GCN5 display increased sensitivity to salt stress because of perturbed H_2_O_2_ production mediated by NADPH oxidases. Wheat GCN5 directly targets gene loci encoding NADPH oxidases leading to their increased H3 acetylation and transcriptional activation in response to salt stress (Zheng *et al*., 2021).

Here, we queried the nuclear sulfenome in *Arabidopsis* by expressing a nuclear version of the YAP1C probe and thereby identified 225 putative redox-active nuclear proteins that undergo sulfenylation. Sulfenylation of the histone acetyltransferase GCN5 at specific cysteine residues was further confirmed by mass spectrometry analysis and their functional significance and impact on protein-protein interaction networks assessed upon cysteine-to-serine mutagenesis.

## Materials and methods

### Plasmid constructs, plant material, and growth conditions

The nuclYAP1C and nuclYAP1A probes were generated by synthesizing the DNA sequence coding for a translational fusion of the following fragments (Supplementary Fig. S1): the Kozak sequence, the nuclear localization sequence (NLS) of the SV40 Large T-antigen (Lee *et al*., 2001), the GSrhino-tag (Van Leene *et al*., 2015) and the codon optimized YAP1C/A sequences, all flanked by the AttL1 and AttL2 Gateway recombination sites. After gene synthesis (Gen9, Cambridge, UK, ceased operations), the sequences were cloned in the plant transformation vector pK7WG2 (Karimi *et al*., 2002) for transformation in cell suspensions. Transgenic PSB-D cultures (NASC stock no. CCL84840) were generated and maintained as described (Van Leene *et al*., 2007). Cells at the mid-log phase (OD_600_ of 0.75) were, unless otherwise specified, treated with 1 mM H_2_O_2_ for 15min. For the subcellular localization experiments, it was cloned in pK7FWG2 for the GFP C-terminal fusion constructs, after omission of the stop codon. T-DNA insertion mutant *gcn5* (SALK_048427) was obtained from the Arabidopsis Biological Resource Center (https://abrc.osu.edu) and verified by genotyping using Phire Plant Direct PCR Master Mix (Thermo Fisher Scientific) with primers listed in Supplementary Table S1. Different transgenic lines expressing GCN5 and cysteine-mutagenized GCN5 in the *gcn5* mutant background were generated using Gateway cloning. To generate the *pGCN5::GCN5-GFP* construct, the *AtGCN5* (*At3g54610*) gene region including introns and its promoter (1.8kb) was amplified from genomic DNA with iProof™ high-fidelity DNA polymerase (Bio-Rad) using attB-flanked primers (Supplementary Table S1). To obtain cysteine-mutagenized GCN5 variants (*pGCN5::GCN5(GCN5C293S)-GFP(gcn5)*, *pGCN5::GCN5(GCN5C368S)-GFP(gcn5)*, and *pGCN5::GCN5(GCN5C400S)-GFP(gcn5)*), point mutations were introduced into the *GCN5* coding sequence to change individual TGT (Cys) or TGC (Cys) codons into TCT (Ser) or TCC (Ser) codons, respectively, using the QuikChange site-directed mutagenesis kit (Agilent Technologies) according to the manufacturer’s instructions. All constructs were sequenced to confirm that no additional mutations had been introduced. PCR products were initially introduced into pDONOR211 entry vectors and then transferred to pB7FWG,0 destination vectors via LR reaction. The final plasmid constructs were heat shocked-transformed in *Agrobacterium tumefaciens* C58C1RifR and introduced into the *gcn5* mutant background by floral dip (Zhang *et al*., 2006). Transgenic lines displaying 3:1 segregation ratio upon spraying with glufosinate-ammonium (560 µl/l) at 5, 9, 11 and 13 days after germination on soil were selected and at least three independent homozygous lines retained in the T3 generation.

To generate transgenic *Arabidopsis* cell cultures expressing N- and C-terminal GFP-tagged GCN5 driven by the cauliflower mosaic virus (CaMV) 35S promoter, PCR products amplified from *Arabidopsis* Col-0 cDNA were first cloned into pDONOR211 vector and then transferred to destination vectors pB7FWG2 (*35S::GCN5-GFP*) and pB7WGF2 (*35S::GFP-GCN5*) via LR reactions. Transgenic cultures were generated by *Agrobacterium* co-cultivation using light-grown *Arabidopsis* PSB-L cell suspension cultures (NASC stock No. CCL84840) as described previously (Forreiter *et al*., 1997). Cultures expressing the bait protein were subcultured in fresh MSMO medium (4.43 g/l Murashige and Skoog basal salts with minimal organics, 30 g/l sucrose, 0.5 mg/l α-naphthaleneacetic acid, 0.05 mg/l kinetin, pH 5.7) at 21°C under 16 h day/8 h night regime with gentle agitation (130 rpm) and subsequently upscaled to 1 liter for GFP pull-down analysis.

For paraquat-induced oxidative stress treatments, seeds were surface-sterilised by fumigation with chlorine (Cl_2_) gas and sown on half-strength Murashige and Skoog (½MS) medium (pH 5.8) containing 0.8% (w/v) sucrose and 30 nM methyl viologen (1,1’-dimethyl-4,4’-bipyridinium dichloride). Seeds were stratified at 4°C in the dark for 2 days and transferred to a growth chamber under long-day conditions (16 h day/8 h night, 21°C).

For in soil growth conditions, the seeds were placed on Jiffy soil peat pellets (44 mm) under a 16 h/8 h day/night regime at 21°C and moderate light intensity conditions (100 µE),

### Protein homology modelling

A three-dimensional model of the AtGCN5 protein was constructed by multi-domain homology modelling using the MODELLER plugin in the Chimera graphical interface (Pettersen *et al*., 2004; Webb and Sali, 2016). Yeast HAT and bromodomain structures were used as separate templates (PDB IDs: 6cw2 and 1e6i). The structure of the linker region was generated separately using the I-TASSER on-line service (Zhang, 2008; Roy *et al*., 2010; Yang *et al*., 2015).

### Recombinant protein expression and purification

The *AtGCN5* coding sequence amplified from *Arabidopsis* Col-0 cDNA using iProof™ High-Fidelity PCR Kit (Bio-Rad) was inserted into pDONR221 vector (Thermo Fisher Scientific) and subsequently subcloned into pDEST-HisMBP (Addgene) expression vector by Gateway cloning. Used primers are listed in Supplementary Table S1. The final hisMBP-GCN5 construct was transformed into *Escherichia coli* strain BL21 (DE3). Protein expression was induced when cell density reached 0.6 (600 nm) by addition of 0.4 mM isopropyl thio-β-D-galactoside for 16 h at 18°C. Cells were harvested by centrifugation and the resulting pellet stored at -20°C until processing. Cell pellet was resuspended in lysis buffer containing 50 mM Tris-Cl (pH 8.0), 200 mM NaCl, 20 mM MgCl_2_, 10% (v/v) glycerol, 0.1 mg/ml 4-(2-aminoethyl) benzenesulfonyl fluoride hydrochloride (AEBSF), 1 µg/ml leupeptin and 0.75 mg/ml lysozyme and incubated on ice for 30 min. After sonication, the lysate was cleared by centrifugation at 18,000*g* for 30 min. The supernatant was incubated with Ni^2+^ Sepharose in a buffer with 50 mM Tris-HCl, (pH 8.0), 200 mM NaCl and 10 mM imidazole for 1 h. Then, the Ni^2+^ Sepharose was packed in an empty PD-10 column and washed with a washing buffer (50 mM Tris-HCl, pH 7.5, 200 mM NaCl and 40 mM imidazole). By using the elution buffer containing 50 mM Tris-Cl (pH 7.5), 200 mM NaCl and 250 mM imidazole, hisMBP-GCN5 protein was eluted. The flow throughs were collected, and the concentrations of purified proteins were measured with the ND-1000 spectrophotometer. The purity of the recombinant protein was assessed by SDS-PAGE and western blotting.

### Transient expression and confocal microscopy

*Agrobacterium tumefaciens* (C58C1; OD_600_ = 0.5, resuspended in 10 mM MgCl_2_, 10 mM MES monohydrate [pH 5.6] and 100 µM acetosyringone) containing p35S::nuclYAP1C-GFP or p35S::nuclYAP1A-GFP were infiltrated with a needleless syringe in the interveinal leaf sections of 6-week-old *Nictiana benthamiana* plants. Three days after infiltration, GFP fusion proteins were visualized. Images were obtained with an inverted Zeiss LSM 710 confocal microscope. Excitation and detection windows were: GFP excitation at 488 nm, emission at 493-543 nm.

### NuclYAP1C/A protein extraction, western blotting and tandem-affinity-purification

Harvested and frozen cells were crushed with pre-cooled pistil and mortars in ice-cold extraction buffer (25 mM Tris-HCl, pH 7.6, 15 mM MgCl_2_, 150 mM NaCl, 15 mM p-nitrophenyl phosphate, 60 mM β-glycerophosphate, 0.1% (v/v) Nonidet P-40, 0.1 mM Na_3_VO_4_, 1 mM NaF, 1 mM phenylmethylsulfonyl fluoride, 1 µM E64, EDTA-free Ultra Complete tablet [Roche; 1/10 mL], 5% (V:V) ethylene glycol, 0.1% (v/v) benzonase, 10 mM iodoacetamide (IAM) and 10 mM N-ethylmaleimide (NEM). Cell debris was removed by two-fold centrifugation at 4°C for 20 min at 20,800*g*. Proteins were separated on a 4% to 12% gradient SDS-PAGE gel and visualized by western blot analysis using 1:5000 diluted peroxidase-anti-peroxidase (PAP; Sigma-Aldrich) antibody-complex. The membrane was visualized using the Western Lightning Plus-EC (Bio-Rad). After Removal of Excess substrate, it was placed into a film cassette, with a chemiluminescence film (Hyperfilm™ EC; GE Healthcare) for approximately 20 s in a dark room. The film was developed with an autoradiogram.

The protocol for affinity purification (Waszczak *et al*., 2014) was modified as follows. Protein extracts were filtered on a glass fiber-pre-filter combined with a 0.45-µm syringe filter (Sartorius Technologies, Germany). Next, 25 mg of protein was incubated with 25 µl IgG-sepharose 6 FAST Flow beads (GE Healthcare) for 1 h on a rotator at 4°C. The beads were washed on Mobicol columns (IMTEC Diagnostics, Belgium) with a 35 µm bottom filter (IMTEC Diagnostics; Belgium) with 150 column volume (CV) of wash buffer (10 mM Tris-HCl, pH 7.6, 150 mM NaCl, 0.1% (v/v) Nonidet P-40, 0.5 mM EDTA, 1 µM E64, 1 mM phenylmethylsulfonyl fluoride, and 5% ethylene glycol) and subsequently incubated for an additional hour with 10 units of PreScission Protease (Sigma-Aldrich; US) at 4°C on a rotator. The elution was collected in a new lowBind Eppendorf tube (Eppendorf; Germany) by gentle spinning at 240*g* and subsequently incubated on 25 µl of Streptavidin Sepharose High Performance (GE Healthcare; US) beads for 1 h at 4°C. Beads from samples from cell suspensions were washed with 100 CV of wash buffer. To elute disulfide-bounded proteins from the YAP1C probe, beads were first incubated with 5 mM dithiothreitol (DTT) for 30 min at room temperature. Elutions were collected by spinning at 1500 rpm. Finally, beads were eluted in a second step with 20 mM desthiobiotin for 5 min at room temperature. NUPAGE Sample buffer (1×) was added to the samples. Gel separation, in-gel trypsin digestion was performed as described (Waszczak *et al*., 2014). Samples were analyzed through liquid chromatography-tandem MS analysis using the LTQ Orbitrap Velos mass spectrometer (Thermo Fisher Scientific) as described by (Waszczak *et al*., 2014). AGC target of 3e6, a max injection time of 20 ms, and a mass range from m/z 300 to 1400. HCD MS/MS spectra were recorded in the data-dependent mode using a Top-20 method with a resolution of 17,500, an AGC target of 1e6, a max injection time of 60 ms, a 1.6 m/z isolation window and normalized collision energy of 30. Peptides m/z that triggered MS/MS scans were dynamically excluded from further MS/MS scans for 18 s.

### Histone acetyltransferase (HAT) activity assay

A colorimetric histone acetyltransferase activity assay kit (ab65352, Abcam; Cambridge, MA, USA) was used to measure *in vitro* HAT activity according to the manufacturer’s instructions. Briefly, 50 µg purified hisMBP-GCN5 protein dissolved in 40 µl water was added to a 96-well plate and mixed with 68 µl of Assay Mix. Absorbance was measured at 440 nm every 30 min and blank values from water-only samples subtracted from their corresponding replicates. HAT activity was expressed as relative units A_440nm_/min/µg of total protein.

### Dimedone labelling and sulfenylation detection by mass spectrometry

A procedure adopted from Huang *et al*. (2018) was used to label recombinant hisMBP-GCN5 with dimedone (5,5-dimethyl-1,3-cyclohexadione). Briefly, purified hisMBP-GCN5 was incubated with 20 mM DTT to reduce all free thiols and excess DTT was removed using Bio-Spin size exclusion columns (Bio-Rad). Next, 12 µM reduced hisMBP-GCN5 protein was incubated with 2 mM dimedone and H_2_O_2_ in different concentration ratios (1:0, 1:5, 1:20, 1:100) for one hour in the dark at room temperature. The excess H_2_O_2_ and dimedone were removed using Bio-Spin size exclusion columns and 4mM N-ethylmaleimide (NEM) was added to the reaction mixture and incubated in the dark at room temperature for 10 min to block any free thiols. Protein samples were separated on 12% non-reducing SDS-PAGE gel and cysteine-dimedone protein adducts visualised by western blot analysis using 1:10,000 diluted rabbit anti-cysteine sulfenic acid primary antibody (Merck Millipore) and 1:10,000 diluted horseradish peroxidase conjugated goat anti-rabbit IgG secondary antibody (Thermo Fisher Scientific) and Western Lightning PlusECL, Enhanced Chemiluminescence Substrate (PerkinElmer).

For mass spectrometry (LC-MS/MS) detection of cysteine-dimedone adducts, protein samples prepared as described above were run on 12% non-reducing SDS-PAGE gel and stained with Coomassie Brilliant Blue (CBB). After destaining, the protein samples were trypsin-digested and the corresponding peptides analysed as described previously (Tossounian *et al*., 2020).

### Immunoprecipitation–mass spectrometry (IP–MS/MS) analysis

IP-MS/MS analysis was performed according to Wendrich *et al*. (2017) with minor modifications. Briefly, three grams of cell culture or plant material were harvested and ground to powder under liquid nitrogen. The resulting powder was resuspended in nine ml of extraction buffer (50 mM Tris-HCl, pH 7.5, 150 mM NaCl, 1% (v/v) detergent Tergitol-type NP-40, Pierce protease inhibitor tablet) and sonicated. The extracts were centrifuged at 18,000*g* (4°C for 15 min), the supernatants passed through a 40-μm cell filter and subsequently incubated with 100 µl anti-GFP magnetic beads (Miltenyi Biotec) at 4°C for one h. The supernatants were transferred to MACS^®^ columns placed in a magnetic rack and the collected beads rinsed four times with 200 µl extraction buffer and two times with 500 µl 50 mM NH_4_HCO_3_. The columns were transferred to protein LoBind tubes and eluted twice with 50 µl pre-heated (95°C) 50 mM NH_4_HCO_3_. An aliquot from the second elution was separated on 12% SDS-PAGE gel and GFP-tagged GCN5 visualised by western blot analysis using anti-GFP antibody followed by anti-mouse horseradish peroxidase (HRP) secondary antibody.

To prepare samples for mass spectrometry analysis, the first eluate was incubated with 1 µl of 500 mM DTT (dithiothreitol) dissolved in 50 mM NH_4_HCO_3_ for 2 h at 60°C. Subsequently, 1 µl of 750 mM iodoacetamide in 50 mM NH_4_HCO_3_ was added and the samples incubated in darkness for 2 h at room temperature. The alkylation was stopped by the addition of 1 µl of 200 mM cysteine and the samples digested overnight with 1 µl of 0.5 µg/μl trypsin (MS Gold, Promega). The resulting peptides were desalted and subsequently analysed on UltiMate™ 3000 RSLCnano System connected to LTQ Orbitrap Velos mass spectrometer (Thermo Fisher Scientific).

### MS/MS data processing and analysis

The resulting raw files were analysed (FDR 1%) in label-free quantification mode using MaxQuant (v1.5.2.8). Spectra were searched against the TAIR10 database and a database containing commonly observed contaminants. Fixed modifications were set to carbamidomethylation, whereas variable modifications were set to oxidation, acetylation and deamidation. Two missed cleavages were allowed for trypsin digestion. Matching between runs was enabled with a matching window time of 2 min. The MaxQuant output was further loaded in Perseus (v1.5.1.6) and proteins identified by only one peptide or no unique peptide were filtered out. Proteins that were enriched at least two times (FC > 2) and had *P* value ≤ 0.05 according to a Student’s *t*-test were considered putative interactors.

### Confocal microscopy

Cell suspensions and plant roots were scanned for GFP signal with a confocal microscope (Zeiss LSM 710, objective: 40× water immersion) using an argon laser (488 nm). The GFP emission was captured between 515 and 545 nm. Auto fluorescence of chlorophyll was monitored in a second channel between 560 and 650 nm. The images were processed using the Zeiss ZEN2009 software.

### Metabolite profiling

For metabolite analysis, 100 mg of homogenised plant material was extracted in a mixture of methyl tert-butyl ether (MTBE):methanol (3:1, v:v) spiked with 25 µg/ml L-Valine-5-13C. Samples were vortexed and again homogenised on RETSCH mill (60 s, 30 Hz, precooled racks), followed by sonication in an ice-cold sonication bath (15 min, 45 kHz) and centrifugation (10 min, 18,000*g*, 4°C). An aliquot of 400 µl of supernatant was transferred into a new 2 ml tube and 300 µl of H_2_O and 100 µl of MeOH were added. Samples were then vortexed and put on RETSCH mill (60 s, 30 Hz, precooled racks), followed by centrifugation (10 min, 18,000*g*, 4°C). Aliquot of 200 µl of the lower phase was transferred into a new 1.5 ml tube, reduced on SpeedVac to dryness and kept at -80°C. Standards of amino acids, acids, sugars, secondary metabolites and others were briefly kept on SpeedVac for 15 min before derivatization. Methoximation reagent of 40 mg methoxyamine hydrochloride in 1 ml pyridine was prepared by sonication followed by heating until dissolved (30°C, 1,000 rpm). Samples were resolved in 20 µl of methoximation reagent and incubated (30°C, 90 min, 800 rpm). Samples were then spiked with 80 µl N-trimethylsilyl-N-methyl trifluoroacetamide (MSTFA) and incubated (37°C, 30 min, 800 rpm), followed by centrifugation (2 min, 14,000*g*). Supernatant of 60 µl was transferred into GC-microvials and closed with caps.

Samples were measured on a Q Exactive GC Orbitrap GC-tandem mass spectrometer and Trace 1300 Gas Chromatograph (Thermo Fisher Scientific). Samples were injected into the machine using the split mode (inlet temperature 250°C, splitless time 0.8 min, purge flow 5.0 mL/min, split flow 6.0 mL/min) onto a TG-5SILMS GC Column (30 m × 0.25 mm × 0.25 µm; Thermo Fisher Scientific), using helium as a carrier gas at a constant flow of 1.2 mL/min. A 28 min gradient (70°C for 5 min followed by 9°C per minute gradient to 320°C, 10 min hold time) was used for metabolite separation, and the electron ionisation mode (electron energy 70 eV, emission current 50 µA, transfer line and ion source temperature 250°C) was used for ionisation. The MS operated in the full scan mode, with 60,000 resolution, scan range of 50-750 m/z, and the automatic maximum allowed injection time with automatic gain control (AGC) was set to 1e6.

### Data processing and analysis

Data were analysed with Compound Discoverer 3.2 (Thermo Fisher Scientific) searched against the GC-Orbitrap Metabolomics, NIST2014, and in-house libraries. Final list of metabolites included only those which passed the identification criteria. Raw files from the mass spectrometer were then uploaded into Skyline (Adams *et al*., 2020) and Compound Discoverer. Peaks in Skyline were manually adjusted for each compound and sample, based on retention time (RT) acquired from Compound Discoverer 3.2 and Xcalibur (Thermo Fisher Scientific), and compared with RT of the standards. Peak areas were then exported and normalised based on the peaks of L-Valine-5-13C and fresh weight. Statistical analysis was performed using MetaboAnalyst 5.0 (Pang *et al*., 2021).

## Results

### Characterising the nuclear sulfenome in *Arabidopsis* cells

To capture sulfenylated proteins in the nucleus, we fused the eight amino acids of the nuclear localization signal (NLS) sequence of the SV40 Large T-antigen to the previously reported YAP1C and YAP1A (without the redox active Cys598) probes, generating *p35S::NLS-YAP1C* and *p35S::NLS-YAP1A*. Both modules also contained a GS^rhino^ tandem affinity purification tag (Van Leene *et al*., 2015) translationally fused to a 85-amino acid fragment (from Asn565 to Asn650) of the C-terminal domain of YAP1 (Fig. 1A) (Waszczak *et al*., 2014; De Smet *et al*., 2019). To validate nuclear targeting, we tested the localization of a C-terminal GFP fusion of both probes (*p35S::NLS-YAP1C-GFP* and *p35S::NLS-YAP1A-GFP*) by *Agrobacterium-*based transient expression in epidermal cells of *Nicotiana benthamiana* leaves. With both constructs, GFP signal was observed in the nucleoplasm and not detected in the nucleoli. Weaker GFP signal was also observed in the cytosol. In contrast, a non-targeted cytoYAP1C-GFP only generated a signal in the cytosol (Fig. 1B).

**Fig. 1.**
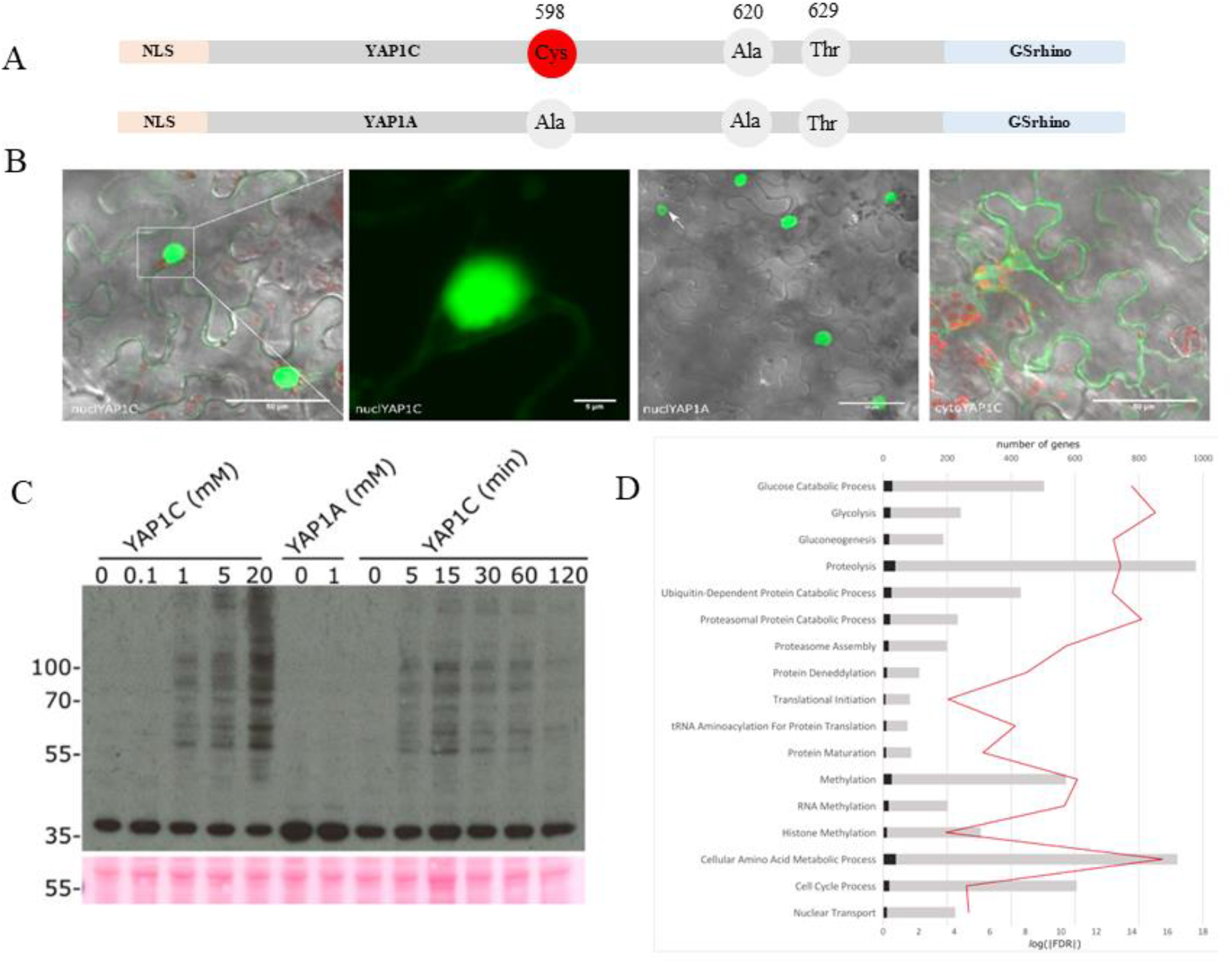
Nuclear sulfenome mining in *Arabidopsis* cells. (A) Schematic depiction of the proteinaceous nuclear sulfenic acid trapping probe YAP1C derived from the yeast AP1-like (YAP1) transcription factor together with the negative control YAP1A lacking a reactive cysteine. (B) Subcellular localization of the nuclear YAP1C probe. Six-week-old *N. benthamiana* plants infiltrated with *p35S::NLS-YAP1C-GFP* (first and second image), *p35S::NLS-YAP1A-GFP* (third image) and non-targeted *p35S:YAP1C-GFP* (fourth image). (C) Time- and dose-response of sulfenic acid formation upon H_2_O_2_ treatment. *Arabidopsis* cells expressing *p35S::NLS-YAP1C-GS* ^rhino^ were treated with 0, 0.1, 1, 5 and 20 mM H_2_O_2_ for 15 min or with 1 mM H_2_O_2_ for 0, 5, 15, 30, 60 or 120 min. As a negative control *Arabidopsis* cells expressing *p35S::NLS-YAP1A-GS* ^rhino^ were treated with 0 and 1 mM H_2_O_2_ for 15 min. Protein bands were visualised with anti-cysteine sulfenic acid antibody. Ponceau staining shows the loading control. NuclYAP1C and nuclYAP1A probes migrate as a single band at ∼35 kDa. (D) Gene Set Enrichment Analysis (GSEA) of proteins found to be sulfenylated after treatment with 1 mM H_2_O_2_ for 15 min using the PlantGSEA toolkit. The identified proteins (black bars) are plotted with the total number of genes for each gene set (grey bar; top axis). False discovery rate (FDR) is shown as a red line (secondary axis at the bottom).

We then evaluated the YAP1C and A protein levels and their capacity to interact with sulfenylated proteins in stable transgenic *Arabidopsis* cell lines expressing GS^rhino^-tagged *p35S::NLS-YAP1C* and *p35S::NLS-YAP1A. Arabidopsis* cells in the mid-log growth phase were exposed to varying concentrations of H_2_O_2_ (0.1, 1, 5, and 20 mM) for 15 min or 1 mM H_2_O_2_ for extended durations (5, 15, 30, 60, and 120 min). Both nuclear YAP1C and YAP1A exhibited a migration pattern as a 35 kDa band, with no high molecular weight bands detected before H_2_O_2_ treatment (Fig. 1C). Formation of complexes with sulfenylated proteins (manifesting as higher molecular weight bands) was observed in a dose-responsive manner only in cells expressing nuclear YAP1C, but not nuclear YAP1A, treated with 1 mM or higher H_2_O_2_ concentrations. Complex formation appeared stable from 5 min to one hour after H_2_O_2_ treatment but diminished noticeably two hours post-treatment. These findings were consistently replicated in three independent experiments. Based on these results, the interactomes of both nuclear YAP1C and YAP1A were isolated through tandem affinity purification from cells treated for 15 min with 1 mM H_2_O_2_ to maximise complex formation. These conditions were identical for the previous identification of the plastid sulfenome (De Smet *et al*., 2019). Proteins identified after tryptic digestion followed by LC-MS/MS present in at least half of the replicates and represented by at least two-high confident peptides, from which at least one being unique, were retained. As such, 375 and 150 proteins were identified to interact with the nuclear YAP1C (Supplementary Dataset S1) and nuclear YAP1A probe (Supplementary Dataset S1), respectively. We considered the nuclear YAP1A interactome as non-specific to the nucleophilic Cys598 of YAP1C and hence retained for further analysis 225 putative sulfenylated proteins. With 85% of these interactors already annotated as nuclear proteins, either through experimentation or prediction based on the presence of a canonical NLS, there was a significant enrichment for nuclear proteins, reflecting an effective targeting of the probe into the nucleus. To identify enriched nuclear processes that are sulfenylated, we performed a gene ontologies (GO) analysis through the PlantGSEA toolkit (Yi *et al*., 2013). Among the enriched GO terms (Fig. 1D) were cell cycle process (19/586; FDR = 2.07E^-5^), nuclear transport (12/213; FDR = 1.25E^-5^), histone methylation (27/545; FDR = 1.17E^-11^) and translational initiation (50/1342; FDR = 1.51E^-17^).

### The histone acetyltransferase GCN5 is sulfenylated at specific cysteine residues

To assess the functional consequences of nuclear cysteine sulfenylation, we focused on the histone acetyltransferase GCN5 that was identified as an interactor of the nuclear YAP1C probe (Supplementary Dataset S1). Oxidative post-translational modifications of epigenetic regulators are emerging as critical means to control the genome-wide distribution of specific histone marks that can ultimately integrate redox signalling mechanisms and chromatin-mediated transcriptional regulation (Bazopoulou *et al*., 2019). *Arabidopsis* GCN5 (*At*GCN5) contains seven cysteine (Cys) residues, four of which are localised in the HAT domain (Cys221, Cys233, Cys293, Cys368), two in the bromodomain (Cys532 and Cys545), and one in the linker (Cys400) (Fig. 2A). Unlike the experimentally solved protein structures of human and yeast GCN5 homologs, including those revealed through cryo-electron microscopy (cryo-EM) as part of the conformational analysis of the human and yeast SAGA complexes, there is currently no experimental structure available for *At*GCN5 (Wang *et al*., 2020; Zhang *et al*., 2022). Therefore, we modelled the three-dimensional structure of *At*GCN5 through multi-domain homology modelling using MODELLER (Webb and Sali, 2016). The N-terminal region of *At*GCN5 is plant-specific and is likely to be highly disordered according to *ab initio* modelling (PHYRE2; Kelley *et al*., 2015). The obtained MODELLER model (Fig. 2B; Supplementary Fig. S2) showed a good overlap with the AlphaFold predicted structure (Jumper *et al*., 2021; Varadi *et al*., 2022) which clearly outlined the unstructured N-terminal region and the HAT and bromodomain structures connected via a linker (Fig. 2C). Based on these predicted protein structures, all cysteine residues found in the HAT domain are likely to be either fully surface-exposed (Cys233 and Cys293) or partially exposed (Cys221 and Cys368). In the bromodomain, Cys532 can be partially exposed within a cleft on the protein surface whereas Cys545 is more likely to be surface-exposed.

**Fig. 2.**
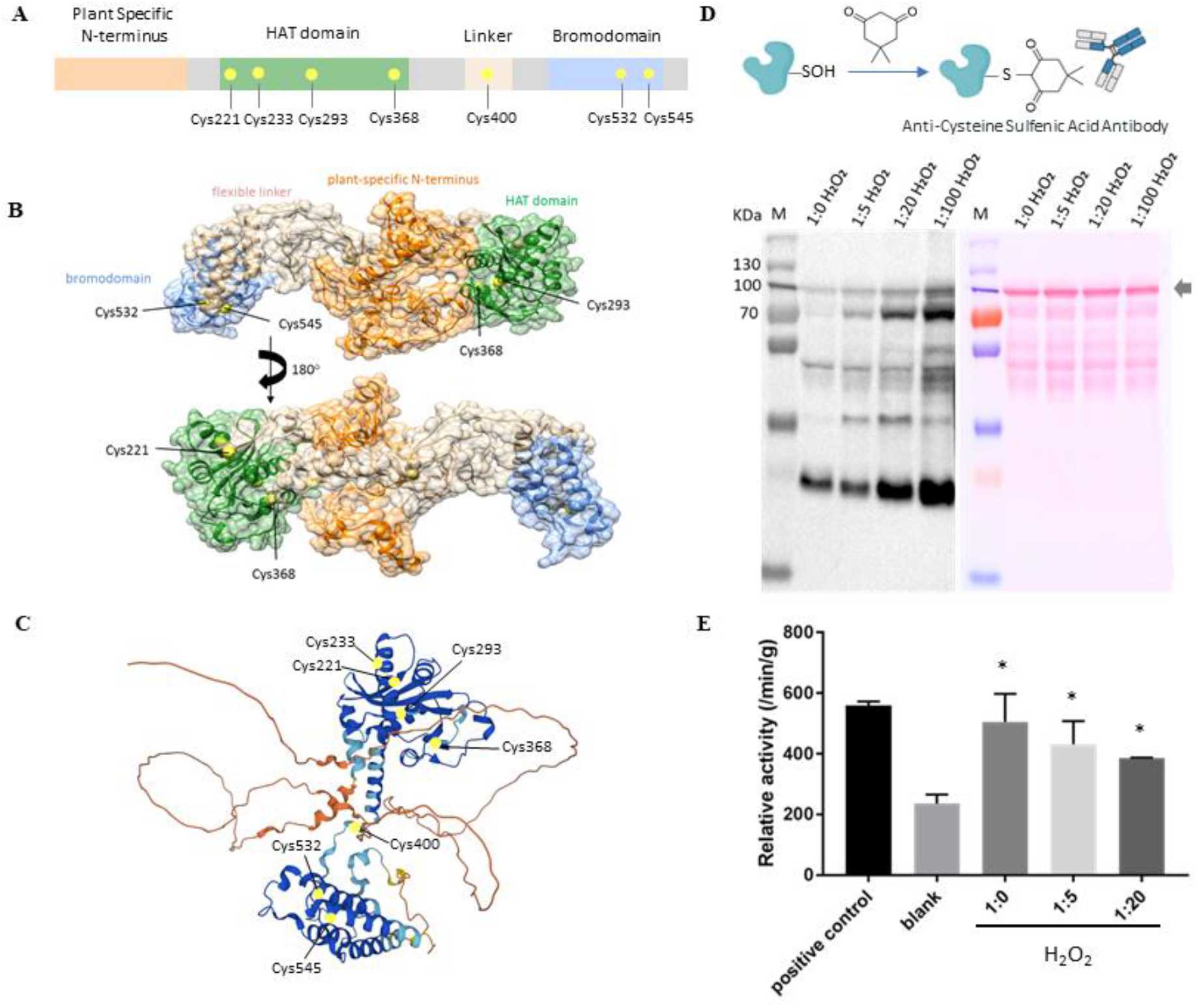
Predicted structure of *Arabidopsis* GCN5 and experimental evidence for its oxidative post-translational modification. (A) Schematic diagram of the GCN5 domains and positions of the individual cysteine residues. (B) Multi-domain homology model of GCN5 according to MODELLER. (C) AlphaFold2 prediction of the GCN5 structure. Coloured based on pLDDT score. (D) Sulfenylation of recombinant His/MBP-tagged GCN5 protein upon H_2_O_2_ treatment. Recombinant GCN5 protein was incubated with increasing H_2_O_2_ concentrations, loaded on a gel and visualised with anti-cysteine sulfenic acid antibody. Ponceau staining shows the loading control. His/MBP-tagged GCN5 migrated as a single band at ∼106 kDa. (E) Histone acetyltransferase activity of recombinant HisMBP-GCN5 in the presence of H_2_O_2._

To assess *At*GCN5 sulfenylation, we first produced recombinant His/MBP-tagged GCN5 protein and reduced it with 20 mM dithiothreitol (DTT). The reduced protein was then incubated with H_2_O_2_, in the presence of dimedone (5,5-dimethyl-1,3-cyclohexadione). Dimedone is a nucleophilic reagent that selectively reacts with the electrophilic sulphur atom in sulfenic acid to form a stable thioether bond (Gupta and Carroll, 2014). The abundance of the resulting cysteine-dimedone protein adducts are first estimated using anti-cysteine sulfenic acid antibodies in a Western Blot analysis. When exposed to varying H_2_O_2_ concentrations ranging from 60 µM to 1.2 mM, the Western Blot bands’ intensities related to His/MBP-GCN5 (106 kDa) exhibited a dose-dependent increase, indicating increased occurrence of cysteine sulfenylation in GCN5 in the presence of H_2_O_2_ (Fig. 2D).

Since four out of the seven cysteine residues present in GCN5 are located in the enzymatic HAT domain, we hypothesised that if one or more of them undergo sulfenylation this is likely to be reflected at the level of histone acetyltransferase activity since oxidative post-translational modifications can alter protein enzymatic activity (Moreno *et al*., 2008). To this end, we monitored the histone acetyltransferase activity of recombinant HisMBP-GCN5 in a fluorescent enzyme coupled-assay using a histone 3-derived peptide as substrate. In the presence of H_2_O_2_, the GCN5 enzymatic activity decreased by 14 and 23% when the molar ratio between the protein and H_2_O_2_ was 1:5 and 1:20, respectively (Fig. 2E).

To pinpoint which cysteine residues undergo sulfenylation when exposed to H_2_O_2_, we excised the protein bands corresponding to sulfenylated HisMBP-GCN5, trypsin digested them and assessed the presence of dimedone adducts on the resulting peptides using liquid chromatography-tandem mass spectrometry (LC-MS/MS). We retrieved 85.7% of the GCN5 sequence and detected all cysteine residues apart from Cys532 and Cys544. Dimedone adducts were identified on peptides containing Cys293, Cys368 and Cys400 (Fig. 3A-C) and their abundance was proportional to the amounts of H_2_O_2_ used for incubation (Fig 3D-F). Taken together, these results show that GCN5 undergoes sulfenylation at Cys293, Cys368, and Cys400 and this negatively affects its enzymatic activity *in vitro*.

**Fig. 3.**
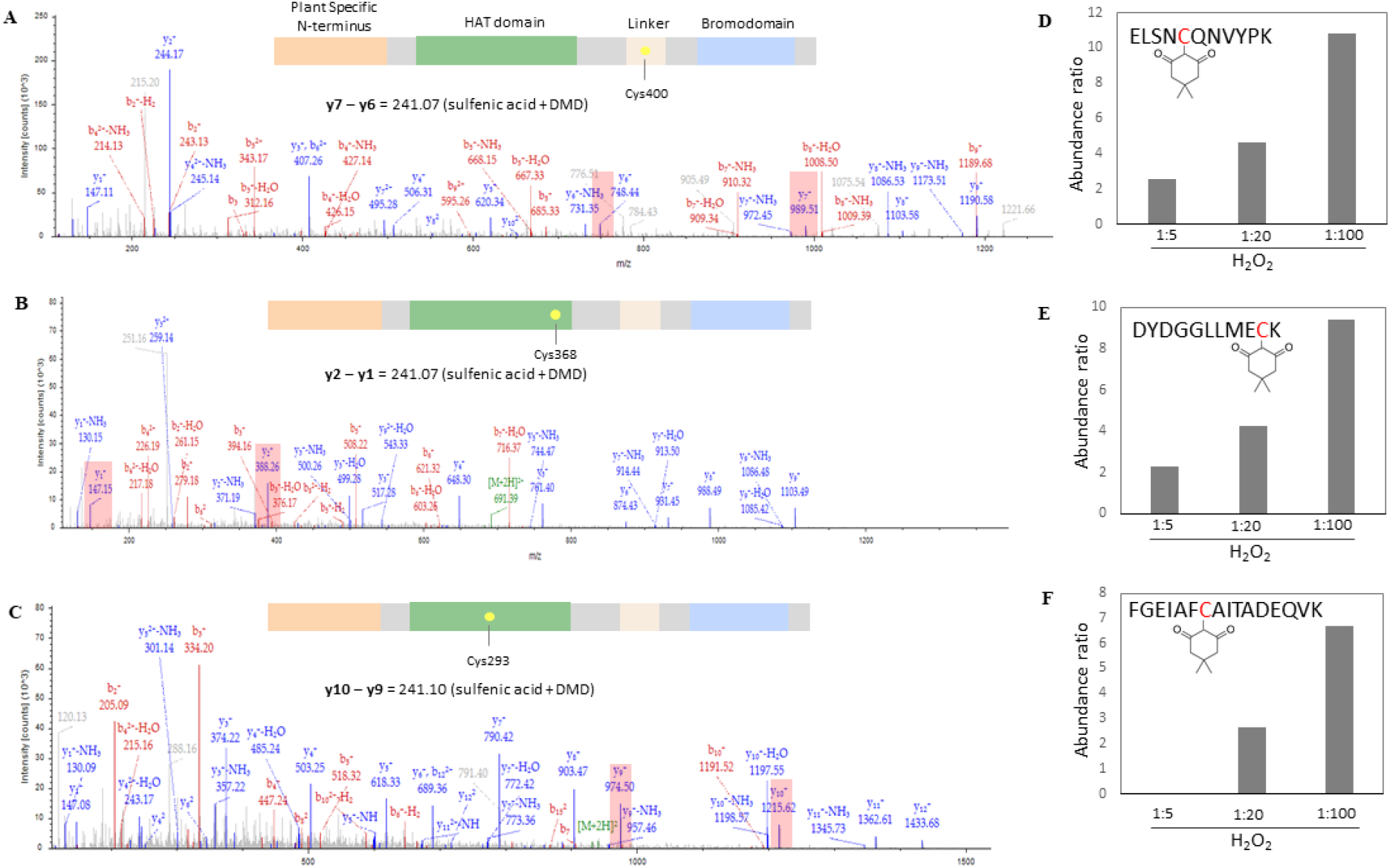
Mass spectrometry-based evidence for GCN5 S-sulfenylation. (A) MS/MS fragmentation pattern of peptide ELSNCQNCYPK containing dimedone-modified sulfenic acid on Cys400. (B) MS/MS fragmentation pattern of peptide DYDGGLLMECK containing dimedone-modified sulfenic acid on Cys368. (C) MS/MS fragmentation pattern of peptide FGEIAFCAITDEQVK containing dimedone-modified sulfenic acid on Cys293. (D) Abundance ratio of label-free quantification (LFQ) intensities of dimedone-modified ELSNCQNCYPK isolated from recombinant HisMBP-GCN5 incubated with varying H_2_O_2_ concentrations relative to H_2_O_2_-free samples. (E) Abundance ratio of label-free quantification (LFQ) intensities of dimedone-modified DYDGGLLMECK isolated from recombinant HisMBP-GCN5 incubated with varying H_2_O_2_ concentrations relative to H_2_O_2_-free samples. (F) Abundance ratio of label-free quantification (LFQ) intensities of dimedone-modified FGEIAFCAITDEQVK isolated from recombinant HisMBP-GCN5 incubated with varying H_2_O_2_ concentrations relative to H_2_O_2_-free samples.

### AtGCN5 cysteine 293 is important for paraquat-induced oxidative stress response

To explore how the redox sensitivity of individual cysteine residues impacts GCN5 function *in planta*, we prepared a panel of transgenic *Arabidopsis* lines in which three cysteine residues (Cys293, Cys368, and Cys400) were mutated to serine and the corresponding constructs driven by the native GCN5 promoter (*pGCN5::GCN5(Cys-Ser)-GFP*) introduced into the *gcn5* mutant background. Independent transgenic lines carrying single cysteine-to-serine mutations and having wild type GCN5 expression levels were selected for homozygosity and evaluated further. *Arabidopsis* mutants lacking functional GCN5 display strong developmental defects such as stunted growth and reduced yield (Vlachonasios *et al*., 2003). Those phenotypes were absent in all complemented *gcn5* lines and they phenocopied the wild type implying that mutagenesis of Cys293 does not affect plant growth and development under control conditions (Fig. 4A).

**Fig. 4.**
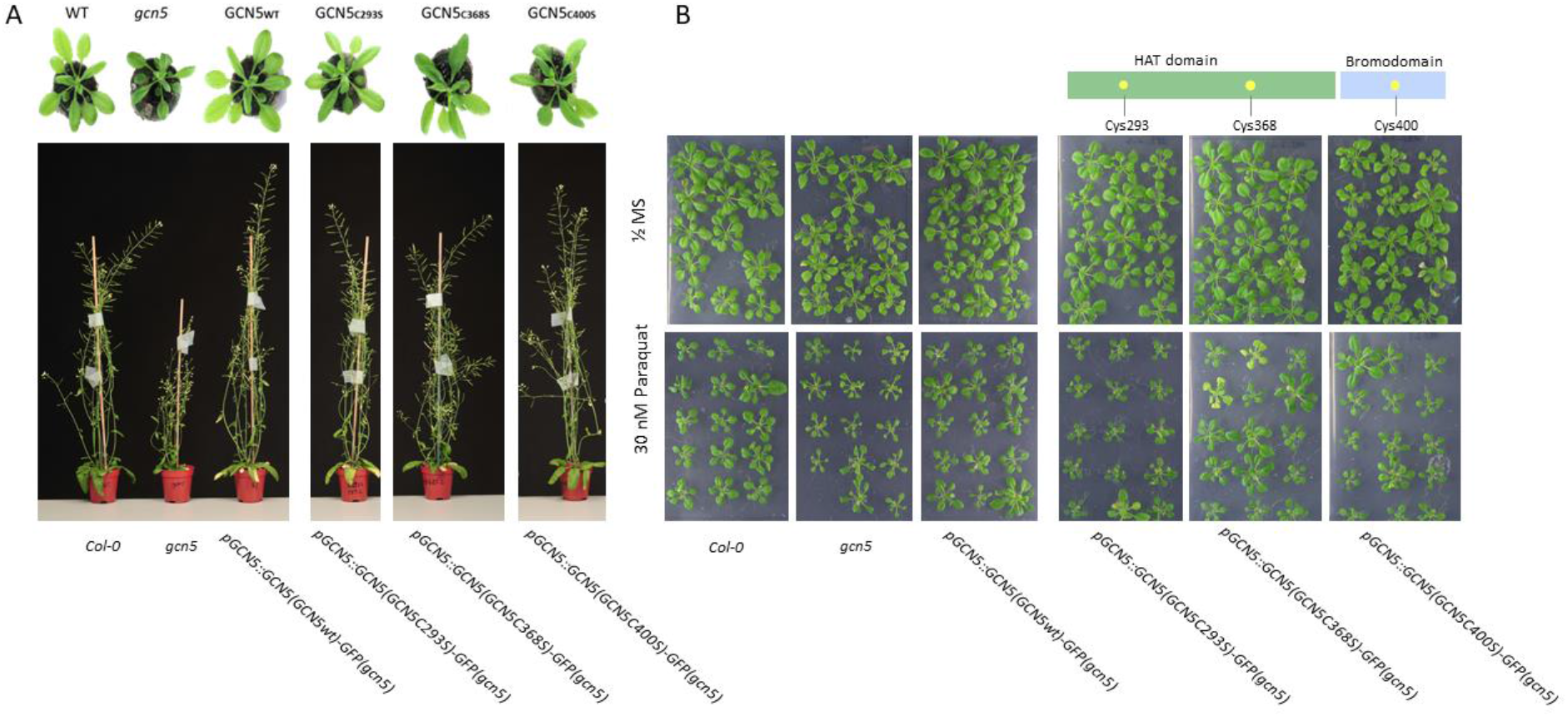
Phenotype of transgenic *Arabidopsis* lines carrying cysteine-to-serine mutated GCN5 variants under control and oxidative stress conditions. (A) Soil-grown wild type (Col-0) and *gcn5* mutant plants together with a panel of complemented *gcn5* mutants carrying cysteine mutagenized GCN5 constructs driven by the native GCN5 promoter. Top panel shows rosettes of 3-week-old plants and bottom panel shows 6-week-old plants. (B) Two-week-old *in vitro* grown plants germinated and grown in the presence of 30 nM paraquat.

In addition to developmental defects, *gcn5* mutants are sensitive to heat and salt stress, both of which are characterised by perturbation of the cellular redox homeostasis (Hu *et al*., 2015; Zheng *et al*., 2019; Sachdev *et al*., 2021). This prompted us to assess the effect of general oxidative stress triggered by the herbicide paraquat (PQ) which generates superoxide at the level of photosystem I (Fuerst and Norman, 1991). The growth inhibition of *gcn5* mutants exposed to 30 nM PQ was more pronounced compared to the wild type (Fig. 4B). Intriguingly, *gcn5* mutant plants complemented with mutated cysteine-to-serine GCN5 variants at positions 368 and 400 were inhibited by PQ similarly to the wild type (Fig. 4B; Supplementary Fig. S3). In contrast, transgenic plants carrying mutagenized Cys293 behaved like *gcn5* mutants (Fig. 4B; Supplementary Fig. S3) and the reduction of rosette area in both genotypes under 30 nM PQ was significantly different than that observed in the wild type (Fig. 5A and B). These results suggest that Cys293 plays an important role during oxidative stress response which could be linked to its sulfenylation under oxidative stimuli.

**Fig. 5.**
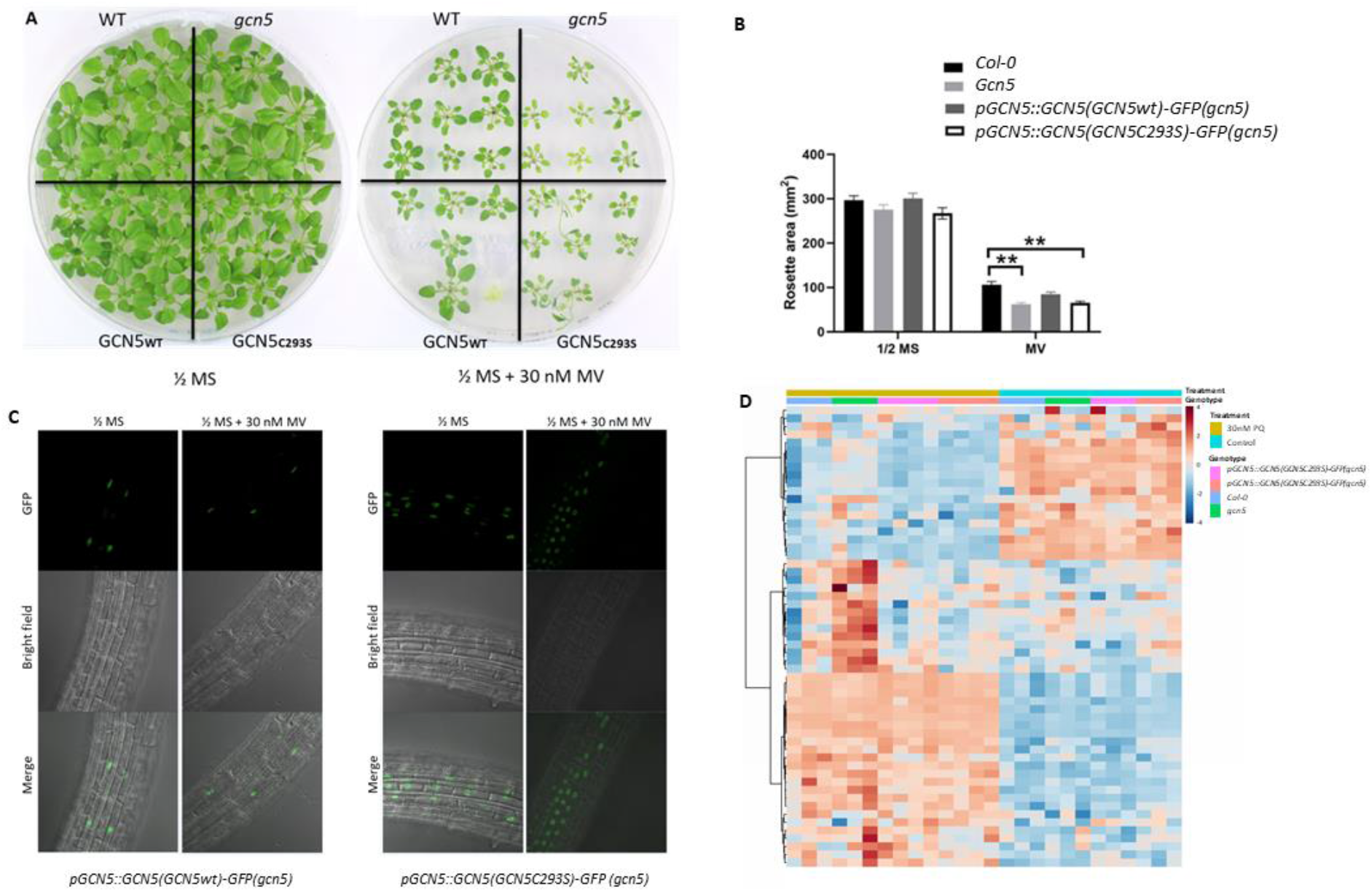
Role of *At*GCN5 Cysteine 293 in paraquat-induced oxidative stress. (A) Three-week-old wild-type (Col-0), *gcn5*, *pGCN5*::*GCN5*(*GCN5wt*)-*GFP(gcn5)*, and *pGCN5*::*GCN5*(*GCN5C293S*)-*GFP(gcn5)* plants germinated and grown on ½MS medium without (left) and with 30 nM paraquat (right). (B) Projected rosette area of 2-week-old *in vitro* grown plants in the presence of 30 nM paraquat. Bars show means of 40 individual rosettes ±SE. Asterisks annotate significant differences according to Student’s *t* test (*P*<0.05). (C) Confocal microscopy images of roots of 7-day-old *pGCN5*::*GCN5*(*GCN5wt*)-*GFP(gcn5)* and *pGCN5*::*GCN5*(*GCN5C293S*)-*GFP(gcn5)* plants grown on ½MS medium without and with 30 nM paraquat. (D) Heat map displaying metabolite abundances in 2-week-old wild-type (Col-0), *gcn5*, and two independent transgenic lines carrying mutagenized Cys293 *pGCN5*::*GCN5*(*GCN5C293S*)-*GFP(gcn5)* germinated and grown on ½MS medium without and with 30 nM paraquat.

Protein S-sulfenylation might have functional roles, such as but not limited to altered protein structure, subcellular localization, and protein-protein interactions (Couturier *et al*., 2013; Huang *et al*., 2018). To explore whether the absence of redox-active cysteine residues impacts the nuclear localisation of *At*GCN5, we visualised the cysteine mutagenized GFP-tagged GCN5 variants and observed nuclear signal under control conditions for all of them (Supplementary Fig. S4). We then probed whether paraquat-induced oxidative stress might lead to a nuclear exclusion of *At*GCN5 using *gcn5* mutants complemented with either *pGCN5*::*GCN5*(*GCN5wt*)-*GFP* or *pGCN5*::*GCN5*(*GCN5C293S*)-*GFP* and observed no changes of the nuclear signal either as a result of paraquat treatment or cysteine mutagenesis (Fig. 5C).

To get an insight into the molecular phenotype that underlies the increased sensitivity of *pGCN5::GCN5(GCN5C293S)-GFP(gcn5)* lines to paraquat, we performed metabolite profiling of primary metabolites using gas chromatography - mass spectrometry (GC-MS) analysis. Under control conditions no statistically significant differences in the levels of the identified metabolites were observed between the wild type, *gcn5* mutants, and Cys293 mutagenized lines (Fig. 5D; Supplemental Dataset S2). Paraquat-induced oxidative stress caused accumulation of glycine, serine, and putrescine in *gcn5* mutant plants in comparison to the wild type, whereas, the levels of those metabolites remained unchanged in two independent transgenic lines carrying mutagenized Cys293 (Fig. 5D; Supplemental Dataset S2).

### Substitution of Cys293 does not dramatically alter the GCN5 protein interactome

AtGCN5 is part of the multiprotein complexes SAGA and PAGA and its sulfenylation might potentially alter the stoichiometry of both complexes. To explore this hypothesis, we first assessed whether using GFP-tagged GCN5 as a bait in an immunoprecipitation assay coupled to mass spectrometry (IP-MS/MS) can inform us on the SAGA composition. Both N- and C-terminal GCN5 GFP fusion constructs driven by the CaMV35S promoter were stably transformed in *Arabidopsis* cell cultures which resulted in a diffused nucleo-cytoplasmic GFP signal (Fig. 6A). Following immunoprecipitation, we identified 35 and 38 significantly enriched proteins in cell lines expressing 35S::GFP-GCN5 and 35S::GCN5-GFP, respectively, which showed a high degree of overlap with 33 proteins were identified in both lines (Fig. 6A, Supplementary Table S2). The identified proteins were highly enriched in SAGA components with most of them displaying various levels of interaction in a STRING analysis (Fig. 6B).

**Fig. 6.**
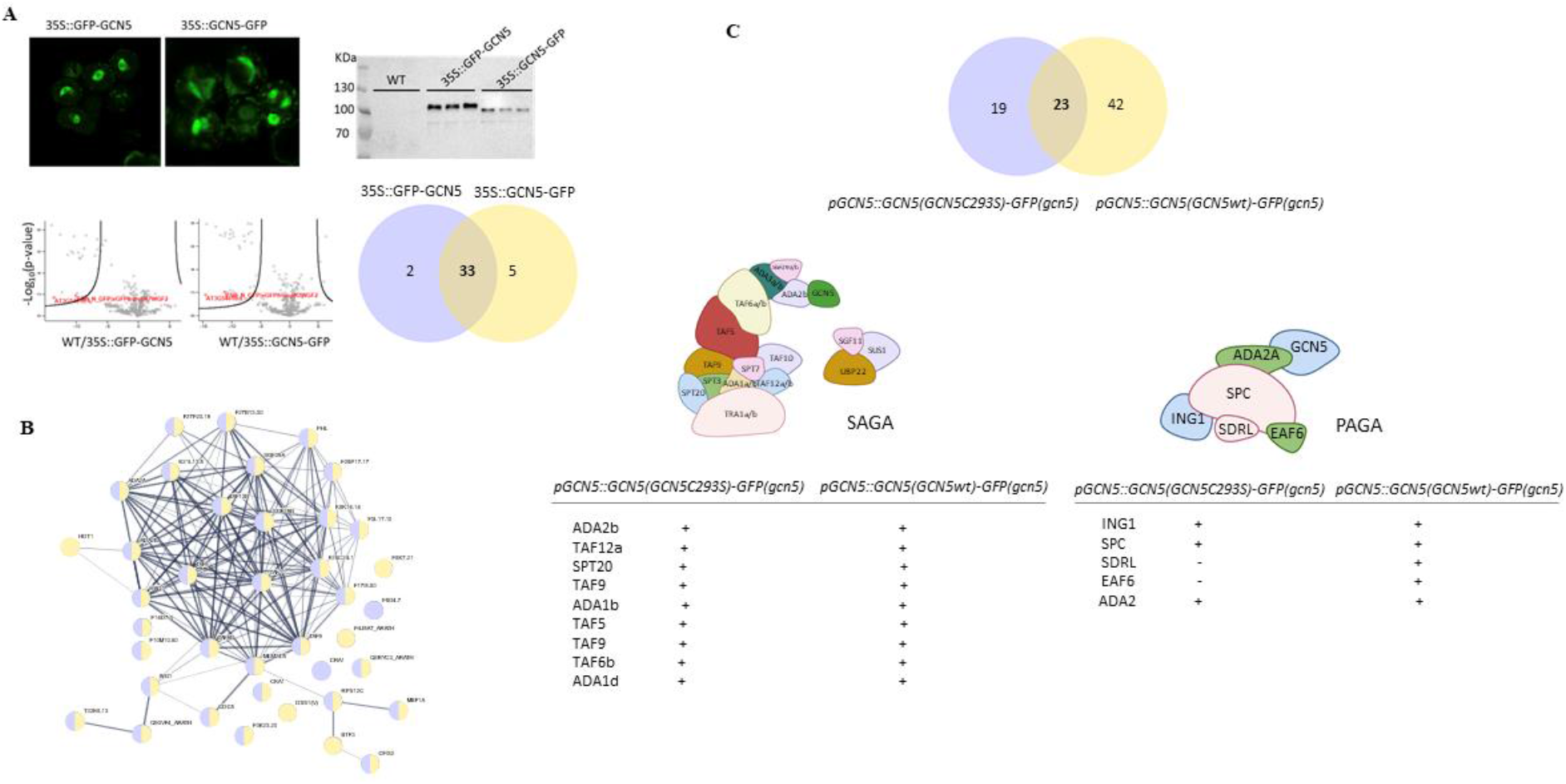
GCN5 protein interactome. (A) Confocal microscopy images of Arabidopsis cell cultures expressing 35S::GFP-GCN5 and 35S::GCN5-GFP together with Western Blot of immunoprecipitated GCN5 visualised with anti-GFP antibody (upper panel). Volcano plot showing differentially abundant proteins that co-precipitated with GCN5 together with a venn diagram showing the overlap between putative protein interactors isolated with N- and C-terminal GFP-tagged GCN5. (B) A protein interaction network generated using STRING showing putative intractors that co-precipitated with N-terminal GFP-tagged GCN5 (yellow) and C-terminal GFP-tagged GCN5 (light violet). C) Venn diagram depicting putative protein interactors that co-precipitated from *pGCN5::GCN5(GCN5wt)-GFP(gcn5)* and *pGCN5::GCN5(GCN5C293S)-GFP(gcn5)* plants together with schematic depiction of SAGA and PAGA complexes. Individual subunits that were isolated in the experiment are listed below and their presence in the individual samples is marked with “+”.

To identify whether part of the GCN5 protein interaction network depends on the presence of redox-active Cys293, we extracted proteins from *Arabidopsis thaliana* seedlings expressing GFP fusions of full-length GCN5 and Cys293S mutated GCN5 in the *gcn5* background to prevent competition with the endogenous bait protein during complex assembly. Nine SAGA subunits co-precipitated with both wild type and Cys293S mutated GCN5 baits (Fig. 6C, Supplementary Table S3). Five components of the PAGA complex were also isolated using wild type GCN5 as a bait, whereas two of them (SDRL and EAF6) were not recovered from *pGCN5*::*GCN5*(*GCN5C293S*)-*GFP(gcn5)* plants (Fig. 6C).

## Discussion

Accumulating evidence reveals that various nuclear proteins, including transcriptional regulators are affected by thiol-based redox mechanisms, which are impacted by the levels of H_2_O_2_ which is one of the major ROS involved in stress signalling cascades operating under adverse environmental conditions. Here, we aimed to comprehensively map out the nuclear proteome that is susceptible to thiol-oxidation. Therefore, we engineered a nuclear localization sequence into a genetically encoded proteinaceous bait that can trap *in cellulo* sulfenylated proteins and expressed it in *Arabidopsis* cells. We report on the identification of 225 sulfenylated nuclear proteins in cultured cells and focus on the functional significance of S-sulfenylation detected on the histone acetyltransferase GCN5.

To identify nuclear proteins that undergo sulfenylation upon disturbance of the cellular redox homeostasis, we introduced H_2_O_2_ in the medium of *Arabidopsis* cell cultures carrying the YAP1C bait. Exogenously applied H_2_O_2_ perturbs the cellular redox homeostasis with extended exposures and high concentrations triggering general oxidative stress. As H_2_O_2_ can diffuse in and out of organelles, like chloroplasts (Mubarakshina *et al*., 2010), and through channels, such as aquaporins (Bienert *et al*., 2007), it is likely that the addition of H_2_O_2_ in the growth medium will lead to nuclear H_2_O_2_ accumulation, at least partially, by H_2_O_2_ diffusion. Additionally, apart from diffusion, it has been reported that ROS are actively produced in plant nuclei by cryptochromes (Jourdan *et al*., 2015; El-Esawi *et al*., 2017). Whether active nuclear ROS production is activated upon exogenous H_2_O_2_ treatment is unknown, but the presence of various enzymatic and non-enzymatic antioxidants in the nucleus together with their dynamic partitioning between the nuclear matrix and the cytosol points toward an orchestrated rearrangement of the nuclear redox homeostasis in response to changes in the environment. For example, H_2_O_2_ accumulates in the nucleus of tobacco leaves exposed to high light. The overexpression of stromal ascorbate peroxidase (APX), leading to reduced chloroplast H_2_O_2_ production, hindered the nuclear H_2_O_2_ accumulation. In contrast, cytosolic APX overexpression had no such effect, underscoring the dependency of high light-induced nuclear H_2_O_2_ accumulation on chloroplast H_2_O_2_ production (Exposito-Rodriguez *et al*., 2017).

From the 225 sulfenylated proteins identified in our study, 186 (83%) had previously been reported in six redox-proteomics studies to undergo reversible cysteine oxidation (Supplementary Table S3) by means of differential labelling of oxidised cysteine enriched by biotin affinity (14 proteins; Wang *et al*., 2012) or by thiopropyl sepharose 6B (64 proteins; Slade *et al*., 2015), using OxiTRAQ (60 and 164 proteins; Liu *et al*., 2014, 2015), the chemical DYn-2 probe (39 proteins; Akter *et al*., 2015) or the cytosolic YAP1C probe (40 proteins; Waszczak *et al*., 2014). The high overlap between different experimental methodologies underscores the reliability of our approach to identify sulfenylated nuclear proteins. The functional significance of S-sulfenylation of all identified proteins, however, awaits experimental validation. Apart from redox-sensitive transcription factors that can serve as direct sensors of redox perturbation, the activity of chromatin modifying enzymes can also be altered by ROS resulting in genome-wide alterations of gene expression. Such evidence is currently predominantly found in animal systems but emerging evidence from plants suggests that such mechanisms are likely present in all life forms given the evolutionary conservation of many epigenetic enzymes. For example, the enzymatic activity of histone methyltransferase MLL1 from *Caenorhabditis elegans* is negatively affected by H_2_О_2_ which is likely caused by cysteine oxidation since the thiol-reducing agent dithiothreitol can restore its function (Bazopoulou *et al*., 2019). MLL1 is part of the evolutionary conserved histone methylation complex COMPASS that deposits H3K4me3 during early development. The levels of this histone mark are depleted by transient ROS increase that accompanies early development and ultimately impacts stress resilience and life span.

GCN5 is part of the evolutionary conserved transcriptional coactivator SAGA that in Arabidopsis regulates thousands of genes linked to developmental and stress response programs (Gan *et al*., 2021; Grasser *et al*., 2021). Within the SAGA complex, GCN5 directly interacts with ADA2B (Wu *et al*., 2021), which was among the most enriched proteins that co-precipitated in our IP-MS/MS analysis. Interestingly, GCN5 also forms another protein complex named PAGA that contains six subunits including four plant-specific members (ING1, SPC, SDRL and EAF6). Even though in the PAGA complex GCN5 directly interacts only with ADA2A (Wu *et al*., 2023), we isolated all PAGA subunits using wild-type GCN5 as a bait. ADA2A similarly co-precipitated with GCN5 lacking cysteine 293, but we were unable to retrieve SDRL and EAF6 using the cysteine mutagenized bait. This is likely due to experimental variation since indirect interactors such as SDRL and EAF6 are unlikely to be affected by the mutation of cysteine 293. Similarly, there were no differences in the SAGA subunits isolated from *pGCN5::GCN5(GCN5wt)-GFP(gcn5)* and *pGCN5::GCN5(GCN5C293S)-GFP(gcn5)* plants, suggesting the entirety of the SAGA complex is not likely to be influenced by cysteine mutagenesis.

Thirty-two protein interactors have been reported so far to interact directly with GCN5 using yeast two-hybrid (Y2H) and *in vitro* protein interaction assays (Stockinger *et al*., 2001; Mao *et al*., 2006; Servet *et al*., 2008). Intriguingly, none of GCN5 protein interactors reported previously were detected using IP-MS/MS in our experiments. This likely reflects the intrinsic properties of the immunoprecipitation method which is more geared towards weak and transient interactions and is usually accompanied with a significant rate of false positive interactions (Wendrich *et al*., 2017). Among the interactors identified by Y2H and confirmed by *in vitro* pull downs was a phosphatase that alters the phosphorylation status of GCN5 and inhibits its enzymatic activity (Servet *et al*., 2008). Interplay of phosphorylation and sulfenylation that can operate together or influence each other *in planta* to fine tune GCN5 activity cannot be excluded. GCN5 has been also shown to interact with the transcription factor EMSY-LIKE 4 involved in plant immunity (Gao *et al*., 2007; Tsuchiya and Eulgem, 2011). How SAGA is targeted to specific genome locations is still a matter of debate and interaction of GCN5 with transcription factors can be a plausible explanation. Moreover, such interactions are unlikely to be limited to GCN5 and other SAGA subunits can be also envisaged to be involved in the process. A comprehensive mapping of the interactomes of all SAGA subunits using a range of methods incl. proximity labelling can offer additional insights. Tissue-specific mechanisms likely contribute to the regulation of specific histone acetylation marks deposition. Exploring the protein interactome through tissue-specific targeting of proximity labeling probes like TurboID holds promise as a valuable research direction. Given the limited differences observed in the protein interactors that co-precipitated with wild type and mutated GCN5, it is likely that the absence of Cysteine 293 does not have a significant impact on the GCN5 interactome but rather modifies its enzymatic activity. This is further corroborated by the fact that its enzymatic activity *in vitro* decreased in the presence of H_2_O_2_. Altered enzymatic activity will ultimately impact the level of histone acetylation marks such as H3K9ac, H3K14ac and H3K27ac, which are deposited by GCN5 (Benhamed *et al*, 2006; Servet *et al*., 2010; Dong *et al*., 2021). The ability of GCN5 to acetylate histones is highly dependent on its proximal proteins, and on its own, GCN5 possesses intrinsically weak activity on histone proteins. The *in vitro* acetylation activity of human GCN5 can be boosted in the presence of ADA2B (Gamper *et al*., 2009). This is achieved by the SANT domain of ADA2 which enhances the binding of GCN5 to its substrate acetyl-CoA (Sun *et al*., 2018). Within the plant-specific protein complex PAGA, GCN5 interacts directly with ADA2A, whereas within SAGA its interaction is with ADA2B. The moderate histone acetylation of genome loci mediated by PAGA in comparison to SAGA (Wu *et al*., 2023) can be a result of fine-tuning of GCN5 activity by the PAGA-specific ADA2A subunit. We did not observe any quantitative differences in the levels of ADA2A and ADA2B that co-precipitated with wild type and Cys293 mutagenized GCN5. IP-MS/MS, however, offers limited quantitative accuracy and it is difficult to speculate whether those results reflect the real stoichiometry of the complex *in planta* and have implications for global acetylation of histone marks. Ultimately, a detailed analysis of the levels of histone acetylation marks deposited by GCN5 and their genome-wide distribution will be essential to understand the precise molecular mechanisms underlying the differential responses of redox-insensitive GCN5 under stress.

Taken together, our results indicate that exogenous H_2_O_2_-induced oxidative stress leads to sulfenylation of a variety of nuclear proteins, indicating a significant impact on nuclear redox homeostasis and multiple nuclear processes. Interestingly, oxidative post-translational modifications of epigenetic modifiers, like histone acetyltransferase GCN5, which we observed to be sulfenylated in our experiments, likely contribute to the fine-tuning of transcriptional reprogramming, particularly at the chromatin level.

## Supplementary data

The following supplementary data are available at JXB online.

Fig. S1. Nucleotide sequence of nuclYAP1C and nuclYAP1A

Fig. S2. Multi-domain homology model of AtGCN5 according to MODELLER

Fig. S3. Phenotyping of *gcn5* plants and plants with mutated GCN5 grown under stress

Fig. S4. Subcellular localization of *At*GCN5 and mutagenized GCN5 variants.

Table S1. Primers sequences

Table S2. Proteins enriched in cell lines expressing 35S::GFP-GCN5 and 35S::GCN5-GFP.

Table S3. Putative protein interactors that co-precipitated from pGCN5::GCN5(GCN5wt)-GFP and pGCN5::GCN5(GCN5C293S)-GFP plants.

Dataset S1. Detailed overview of putative sulfenylated proteins identified with nuclYAP1C from *Arabidopsis* cell cultures

Dataset S2. Metabolites

## Author contributions

JM, PIK and F.V.B.: conceptualization; BDS, and DV: methodology; BDS, XY, ZP, AFF, AM, DV, SPdR, and PIK: formal analysis; BDS, XY, ZP, AFF, AD, AM, DV, SPdR, and PIK: investigation; DV, and SpdR: resources; SPdR: data curation; BDS, CC, FVB, and PIK: writing - original draft; BDS, CC, DV, PIK, and FVB: writing - review & editing; AFF: visualization; D.V., JM, and FVB: supervision; BDS, DV, KXC, JM and FVB: funding acquisition.

## Conflict of interest

None to declare.

## Funding

This research was financially supported by the Czech Science Foundation (grant No. 22-17092S) to P.I.K, a personal PhD grant to B.D.S. (Strategisch Basisonderzoek) from IWT-Vlaanderen (project number: 141007), FWO Nucleox grant No G007723N, EOS grant to D.V. and S.P.d.R (No. 30829584), and a FWO postdoctoral fellowship to K.X.C. (No. 12N4818N).

## Acknowledgments

The authors gratefully acknowledge Mihail Angelov for assistance with Dataset S2, Robin Pottie and Kristof Verleye for technical assistance and supervision, and Dr. Martine De Cock for help in preparing the manuscrip.

## Notes

### Competing Interest Statement

The authors have declared no competing interest.

